# Impact of parasitic lifestyle and different types of centromere organization on chromosome and genome evolution in the plant genus *Cuscuta*

**DOI:** 10.1101/2020.07.03.186437

**Authors:** Pavel Neumann, Ludmila Oliveira, Jana Čížková, Tae-Soo Jang, Sonja Klemme, Petr Novák, Katarzyna Stelmach, Andrea Koblížková, Jaroslav Doležel, Jiří Macas

## Abstract

- The parasitic genus *Cuscuta* (Convolvulaceae) is exceptional among plants with respect to centromere organization, including both monocentric and holocentric chromosomes, and substantial variation in genome size and chromosome number. We investigated 12 species representing the diversity of the genus in a phylogenetic context to reveal the molecular and evolutionary processes leading to diversification of their genomes.
- We measured genome sizes and investigated karyotypes and centromere organization using molecular cytogenetic techniques. We also performed low-pass whole genome sequencing and comparative analysis of repetitive DNA composition.
- A remarkable 102-fold variation in genome sizes (342–34,734 Mbp/1C) was detected for monocentric *Cuscuta* species, while genomes of holocentric species were of moderate sizes (533–1,545 Mbp/1C). The genome size variation was primarily driven by the differential accumulation of repetitive sequences. The transition to holocentric chromosomes in the subgenus *Cuscuta* was associated with loss of histone H2A phosphorylation and elimination of centromeric retrotransposons. In addition, the basic chromosome number (x) decreased from 15 to 7, presumably due to chromosome fusions.
- We demonstrated that the transition to holocentricity in *Cuscuta* was accompanied by significant changes in epigenetic marks, chromosome number and the repetitive DNA sequence composition.

## Introduction

Nuclear genomes of flowering plants (angiosperms) show remarkable variation in basic properties, including size, sequence composition, and chromosome organization. Genome sizes span three orders of magnitude, from 0.06 Gbp/1C in *Genlisea tuberosa* (Fleischmann *et al.*, 2014) to 148.8 Gbp/1C in *Paris japonica* (Pellicer *et al.*, 2010); however, most species have relatively small genomes and the giant ones are restricted to a few taxa (Pellicer *et al.*, 2018; Pellicer & Leitch, 2020). Genome duplication and the differential accumulation of repetitive DNA are recognized as the main drivers of genome size diversification (Bennetzen & Wang, 2014; Macas *et al.*, 2015; Kelly *et al.*, 2015). However, the genetic, epigenetic, and ecological factors that determine evolutionary trajectories leading to genomes of various sizes are yet to be fully elucidated. There is also considerable variation in angiosperm karyotypes, with two to over a thousand chromosomes and frequent changes in chromosome number and morphology (Love, 1976; Heslop-Harrison & Schwarzacher, 2011). Perhaps the most dramatic change is the switch between different types of centromere organization. Monocentric chromosomes have centromeres confined to a single specific region observed as a primary constriction during mitosis, while holocentric chromosomes are characterized by centromeres dispersed along the entire chromosome length (Wanner *et al.*, 2015). Species with monocentric chromosomes are more common and supposedly ancestral, and there are several phylogenetic lineages that underwent independent switches to holocentricity (Melters *et al.*, 2012). However, the factors that triggered these transitions and how they impacted other genome features are not known.

To tackle the questions outlined above, comparative analyses of related taxa showing trait variation are often performed. For example, groups of repeats with a high impact on genome size evolution in several plant taxa have been identified by analyzing the repeat composition of individual genomes and evaluating these data with respect to the species phylogeny (Novák *et al.*, 2014; Macas *et al.*, 2015; Kelly *et al.*, 2015; Gaiero *et al.*, 2019). This approach is highly informative when applied to a group of closely-related species (e.g., within a genus) due to their recent ancestry and thus common genetic background. However, these species usually do not vary substantially in the features of interest. Genome sizes of most plant genera vary less than 10-fold, while a 100-fold variation has only been described for much broader taxonomic levels, like tribes or families (Pellicer *et al.*, 2014; Pellicer & Leitch, 2020). Similarly, a holocentric chromosome organization is usually shared within whole families, like Cyperaceae (Cuacos *et al.*, 2015), and thus the closest related taxa possessing monocentric chromosomes are phylogenetically distant.

The genus *Cuscuta* (Convolvulaceae) includes about 200 species of parasitic plants (García *et al.*, 2014) and is exceptional among angiosperm genera with respect to variation in both genome size and chromosome organization. *Cuscuta* species inhabit diverse habitats and hosts on all continents except Antarctica, and are classified into three subgenera, *Monogynella, Grammica,* and *Cuscuta* (García *et al.*, 2014). Holoploid genome sizes (1C) ranging from 0.47 Gbp in *C. campestris* to 32.12 Gbp in *C. indecora* have been reported (McNeal *et al.*, 2007a; Kubešová *et al.*, 2010). Since genome size has only been measured in 15 *Cuscuta* species, it is possible that the variation is even greater. Moreover, species within the subgenus *Cuscuta* possess holocentric chromosomes, while species in the two remaining subgenera include only monocentric chromosomes (Pazy & Plitmann, 1991, 1994, 1995, 2002). Thus, *Cuscuta* is the only well-established case of a switch to holocentrism within the evolutionary history of a single genus.

All *Cuscuta* species are obligate parasites utilizing various host plants, and ecological and physiological adaptations associated with their parasitic lifestyle have been the subject of extensive research. Various metabolites and nucleic acids are exchanged between *Cuscuta* and its hosts (reviewed in (Kim & Westwood, 2015)). Moreover, 108 genic and 42 non-genic DNA regions in *C. campestris* and related species were acquired via horizontal gene transfer (Vogel *et al.*, 2018; Yang *et al.*, 2019). This observation suggests that genome expansion in the genus occurred by the proliferation of horizontally transferred mobile elements. However, studies focusing on repeat composition with respect to species phylogeny and genome size variation in *Cuscuta* are lacking. Genome assemblies of two *Grammica* species, *C. campestris* and *C. australis*, are available; however, both genomes are small and thus repeat-poor, and investigations have emphasized genes (Sun *et al.*, 2018; Vogel *et al.*, 2018). Although relatively few species have been analyzed, it is evident that *Cuscuta* species are extremely variable in chromosome size (0.4–23 μm), number (2n = 8–60), and morphology (Pazy & Plitmann, 1995). The holocentric nature of chromosomes has been demonstrated in all species of the subgenus *Cuscuta* tested thus far (Pazy & Plitmann, 1987, 1991, 1994, 1995; García, 2001; Guerra & García, 2004; Oliveira *et al.*, 2020). By contrast, monocentric chromosomes have been documented in species of the subgenera *Monogynella* and *Grammica* (Pazy & Plitmann, 1995; Oliveira *et al.*, 2020). Although it is assumed that all species in these two subgenera and in the subgenus *Cuscuta* have monocentric and holocentric chromosomes, respectively, the existence of both types of centromere organization in species of the same subgenus cannot be fully ruled out because the majority of *Cuscuta* species have not yet been analyzed. However, the determination of centromere organization in some species is hampered by the small size of chromosomes, which prevents assessments of morphology and behavior during cell division. In addition, while centromere-specific variant of histone H3, referred to as CENH3, appears to be a marker for centromere position in monocentric *Cuscuta* species, it cannot be applied to holocentric *Cuscuta* species, in which the role of CENH3 in centromere determination and function has been questioned (Oliveira *et al.*, 2020).

In this study, we explored the wide variation in genome sizes within different *Cuscuta* lineages, and correlations with the content and composition of repetitive DNA and ploidy levels. We focused on comparisons among species with different types of centromere organization, as determined unambiguously by a combination of conventional cytogenetic methods and the immunodetection of centromeric proteins.

## Materials and Methods

### Plant material

Information on the origin of the plant material and cultivation is provided in Notes S1. *Cuscuta* seeds were germinated as described previously (Oliveira *et al.*, 2020) and cultivated on the following host plant species: *Urtica dioica (C. europaea), Betonica officinalis (C. epithymum), Linum usitatissimum (C. epilinum), Pelargonium zonale (C. japonica, C. monogyna* and *C. reflexa)* and *Ocimum basilicum* (remaining *Cuscuta* species). To prevent the accidental spread of *Cuscuta* species, all plants were grown in isolation and autoclaved when they were no longer needed for experiments.

### Estimation of genome size and analysis of DNA sequences

Nuclear genome size was estimated as described previously (Doležel *et al.*, 2007) and detailed in Methods S1. Genomic DNA was extracted from young shoots as described previously (Dellaporta *et al.*, 1983). Shotgun sequencing of DNA was performed by the University of Rochester Genomics Research Center (New York, NY, USA) and Admera Health (South Plainfield, NJ, USA), employing the Illumina platform to generate 100–150 bp paired-end reads from ~300–500 bp fragment libraries. The sequence data for all twelve species were deposited in the Sequence Read Archive (study accession: ERP117356, Table S1). Before analyses, the raw Illumina reads in FASTQ format were pre-processed using the “Preprocessing of FASTQ paired-end reads” tool (https://repeatexplorer-elixir.cerit-sc.cz/galaxy) with default settings, except all reads were trimmed to 100 nt and reads shorter than 100 nt were discarded.

Repetitive sequences were identified by similarity-based clustering of Illumina paired-end reads using the RepeatExplorer and TAREAN pipelines at https://repeatexplorer-elixir.cerit-sc.cz/galaxy/ (Novák *et al.*, 2013, 2017). The numbers of reads analyzed for individual species are shown in Table S1. All clusters representing at least 0.01% of the genome were manually checked, and the automated annotation was corrected if needed. Finally, the clusters were used to characterize and quantify the most abundant repeats. Genomic proportions of the major repeat types (Table S2) were calculated based on the proportion of reads in individual annotated clusters. To calculate the total proportion of repeats with more than 20 copies in the genome, a random sample of forward reads free of organellar sequences and representing 5% coverage of the nuclear genome was used for an all-to-all similarity search using the optimized BLASTn program mgblast, as implemented in the TGI Clustering Tool (https://sourceforge.net/projects/tgicl/version2.1-1). The following mgblast command line options were used: -W18 -UT -X40 -KT -JF -F “mD” -v100000000 -b100000000 -D4 -C50 – H30. All similarity hits with sequence identities above 80% over at least 55 bp were recorded, and reads with at least one hit were considered repeats with more than 20 copies in the genome. Comparative clustering analysis was performed using RepeatExplorer run in comparative mode using ca 430,000 reads from each species (the maximum number of reads that could be analyzed on the server). Abundances of simple sequence repeats were calculated using Tandem Repeats Finder (TRF; (Benson, 1999)) and TRAP (Sobreira *et al.*, 2006). The input for TRF was prepared by concatenating one million randomly selected reads, each of which was separated by a stretch of 50 Ns. Identification and classification of transposable element protein domain sequences were performed using the DANTE tool at https://repeatexplorer-elixir.cerit-sc.cz/galaxy/ (Novák *et al.*, 2019) and the REXdb database (Neumann *et al.*, 2019).

Confirmation of species identities and phylogenetic relationships were performed using the internal transcribed spacer (ITS) region of 45S rDNA and a gene encoding the large subunit of ribulose bisphosphate carboxylase (*rbcL*). Consensus ITS sequences were extracted from RepeatExplorer clusters. Sequences of *rbcL* were selectively assembled using GRABb (Brankovics *et al.*, 2016) employing as inputs Illumina paired-end reads and a bait file containing a set of *Cuscuta rbcL* sequences downloaded from GenBank. Illumina paired-end reads from *Ipomoea* species were downloaded from SRA (run accessions DRR013919 and ERR1512971). Sequences of ITS and *rbcL* and results of BLASTN searches against GenBank are provided in Tables S3 and S4. Most BLASTN hits were identical or nearly identical sequences from expected *Cuscuta* species, confirming their identity. In some cases, similarly strong hits were obtained for additional *Cuscuta* species, reflecting the close phylogenetic relationships or suggesting that species of some GenBank accessions were misidentified.

A phylogenetic analysis was performed using the neighbor-joining algorithm implemented in SeaView (Gouy *et al.*, 2010). Bootstrap values were calculated from 5,000 replicates. The sequences were aligned using Muscle (Edgar, 2004). Divergence times were estimated using the RelTime method implemented in MEGA X (Mello, 2018) employing trees calculated using PhyML 3.0 with smart model selection (Guindon *et al.*, 2010; Lefort *et al.*, 2017) and assuming that the time to the most recent common ancestor of *Cuscuta* and *Ipomoea* was 33.3 million years ago (Ma; (Sun *et al.*, 2018)). Phylogenetic trees were drawn and edited using Dendroscope (Huson & Scornavacca, 2012). Putative horizontally transferred LTR retrotransposons were identified as described in Methods S2.

### Cytogenetic methods

Chromosome preparation, immunodetection, probe labeling, and fluorescence *in situ* hybridization were performed as described previously (Oliveira *et al.*, 2020). Antibodies to H3S10ph and H2AT120ph were purchased from Millipore (Cat. # 04-1093; Temecula, CA, USA) and MYBioSource (MBS852710; San Diego, CA, USA), respectively. An antibody to CENH3 was custom-produced by GenScript (Piscataway, NJ, USA) against a peptide designed based on the *C. campestris* CENH3 sequence (Oliveira *et al.*, 2020). Sequences of fluorescence *in situ* hybridization (FISH) probes and hybridization and washing conditions are provided in Table S5.

## Results

### Species with holocentric chromosomes form a monophyletic lineage within the genus

Twelve *Cuscuta* species representing all three subgenera as well as different clades defined within the most diverse subgenus *Grammica* were included (Stefanović *et al.*, 2007; García *et al.*, 2014). Due to the poor availability of species from the subgenus *Cuscuta* that were previously reported to possess holocentric chromosomes (Pazy & Plitmann, 1991, 1994, 1995), we selected *C. europaea*, *C. epithymum* and *C. epilinum* as other representatives for this subgenus. To verify the centromere organization in these species, we inspected the chromosome morphology, behavior of chromosomes during mitosis, and distribution of epigenetic histone marks known to occur at centromeric regions during mitosis. Holocentric chromosomes were confirmed for all three species in the subgenus *Cuscuta*, as demonstrated by the absence of primary constrictions, typical chromosome arrangement during mitotic anaphase with longitudinal axes oriented perpendicularly to the axis of division, and chromosome-wide distribution of histone H3 phosphorylated at serine 10 (H3S10ph) (Figs 1, 2, S1). Strikingly, the phosphorylation of threonine 120 of histone H2A (H2AT120ph), previously established as a universal centromere marker in plants (Demidov *et al.*, 2014), was not detected in any of the holocentric species, suggesting that this type of phosphorylation was lost in the subgenus *Cuscuta*. By contrast, chromosomes in species of the other two subgenera possessed primary constrictions and/or displayed the expected single domain localization of both H3S10ph and H2AT120ph as well as the foundational kinetochore protein CENH3 (Fig. S1).

**Figure 1.**
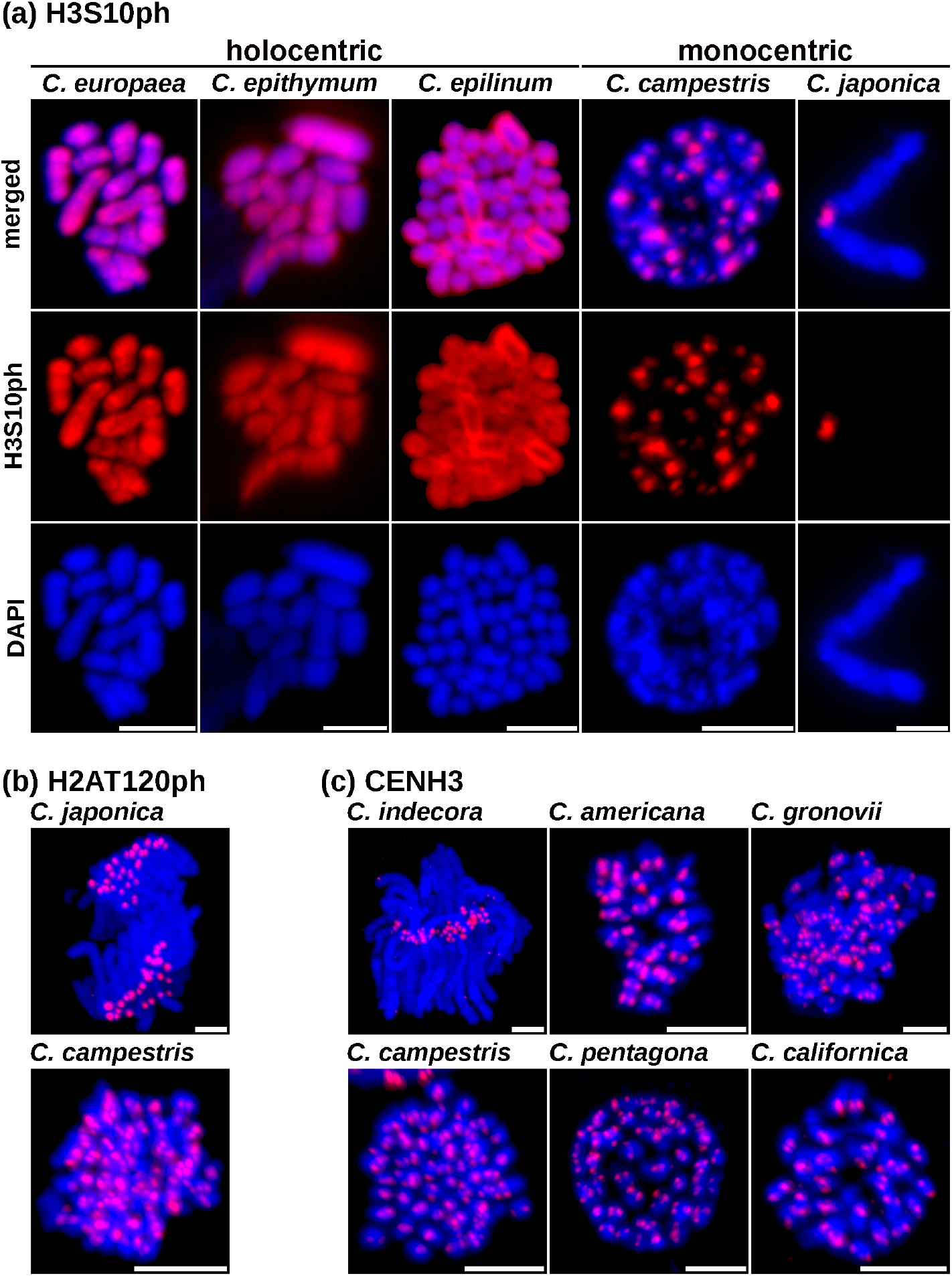
Distribution of centromere markers on mitotic chromosomes of holocentric and monocentric *Cuscuta* species. **(a)** H3S10ph was detected over entire chromosomes in all three holocentric species and at discrete sites in monocentric *Cuscuta* species. Chromosomes of *C. campestris* were too small to exhibit primary constriction, but each displayed a single region with H3S10ph signals, in agreement with the presence of a single centromere domain per chromosome. In *C. japonica,* H3S10ph was confined to primary constriction, demonstrating that H3S10ph is a centromere mark in *Cuscuta* species. **(b)** Detection of H2AT120ph in *C. japonica* and *C. campestris.* **(c)** Detection of CENH3 in all six *Grammica* species evaluated in this study. Chromosomes were counterstained with DAPI (blue). All scale bars = 5 μm.

Phylogenetic trees of the selected species were inferred using sequences of ITS of 45S rDNA and a plastid gene encoding the large subunit of ribulose bisphosphate carboxylase (*rbcL*) (Tables S4, S5). The trees constructed using the neighbor-joining method were supported by high bootstrap values, and the topologies were in complete agreement with each other (Fig. S2a) and with those of previously published trees (McNeal *et al.*, 2007a; García *et al.*, 2014). The phylogenetic analysis showed the basal position of the subgenus *Monogynella* and confirmed that the switch to a holocentric chromosome organization occurred late in the evolution of the genus, along with the diversification of the other two lineages, the holocentric subgenus *Cuscuta* and monocentric *Grammica* (Fig. S2a). Considering that the *Cuscuta* and *Ipomoea* lineages split ~33 Ma (Sun *et al.*, 2018), we estimated that the ancestor of *Monogynella* diverged 23–20.5 Ma and the two sister subgenera *Cuscuta* and *Grammica* split 17.5–17 Ma (Fig. S2b).

### Nuclear DNA amounts vary widely and genome expansion is confined to monocentric lineages

Although genome size estimates were available for some species, we estimated nuclear DNA amounts in all twelve species using the same methodology to enable interspecific comparisons of genome size data. A flow cytometric analysis of nuclei isolated from vegetative shoots revealed substantial differences in holoploid genome sizes (1C). Genomes ranged from 386 Mbp for *C. californica* to 34,734 Mbp for *C. reflexa.* (Fig. 2, Table S6). The genome of *C. australis* (subgenus *Grammica*) was previously reported to be 272 Mbp based on the k-mer distribution (Sun et al., 2018). Using our method, however, we detected a genome size of 342 Mbp. The largest genomes were recorded in all three species of the subgenus *Monogynella* (25,584–34,734 Mbp) and in *C. indecora* (22,675 Mbp) from the subgenus *Grammica*. Other *Grammica* species and all three species of the holocentric subgenus *Cuscuta* had much smaller genomes (342–3579 Mbp and 533–1545 Mbp, respectively) (Fig. 2, Table S6). We detected 102-fold variation in genome size in *Cuscuta* species; in comparison with genome size data from the C-value database (release 7.1, Apr 2019; (Pellicer & Leitch, 2020)), this variation is extraordinary for a single genus and is more similar to genome size variation in entire families (Fig. S3).

**Figure 2.**
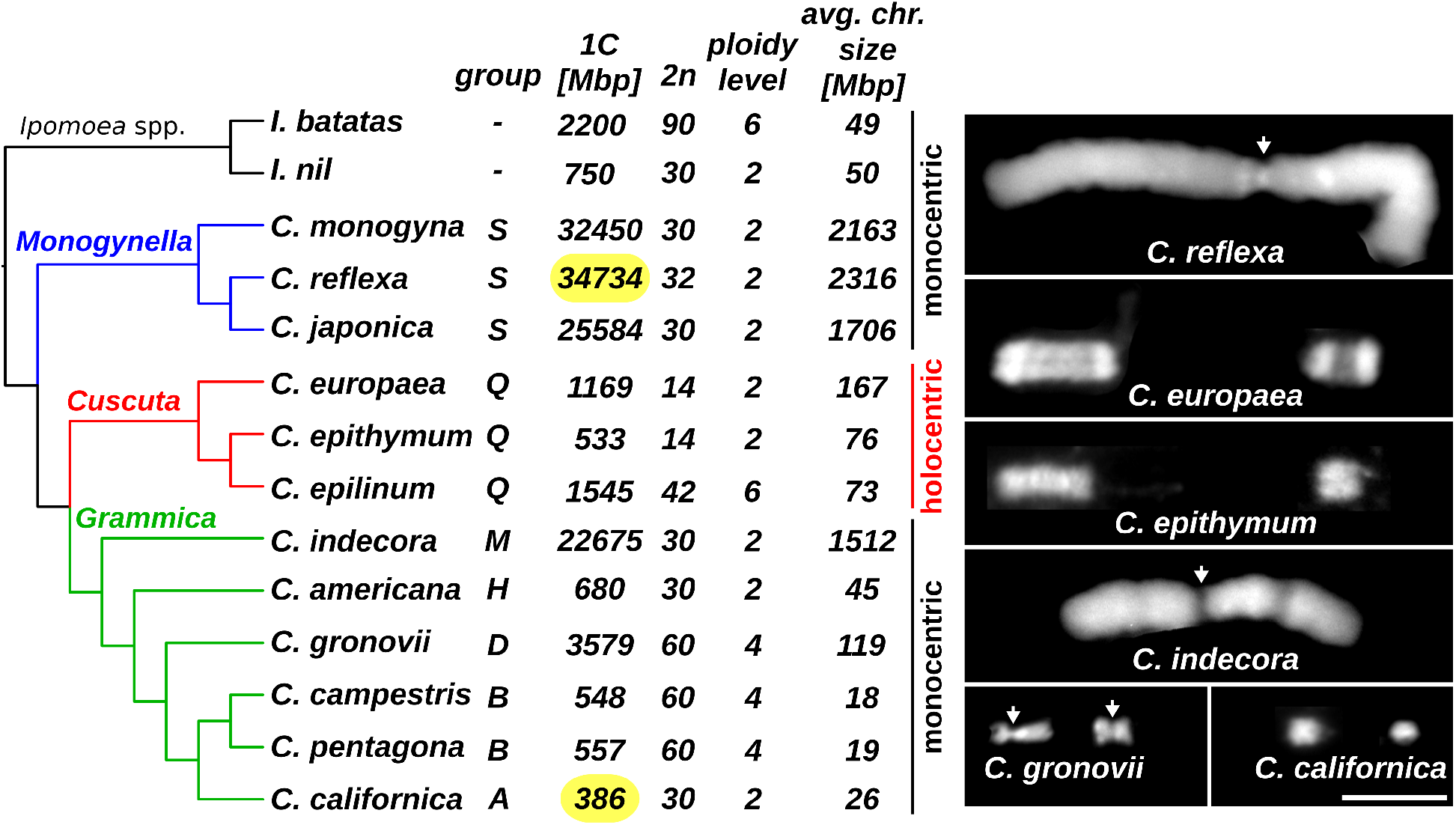
Genome size, chromosome number, and centromere organization in a phylogenetic context. The phylogenetic tree was inferred from ITS and *rbcL* sequence alignments (Fig S2). The classification of *Cuscuta* species to three subgenera and seven groups is based on previous results (García *et al.*, 2014). The genome sizes of all *Cuscuta* species were estimated in this study (Table S5). The smallest and largest genomes of the twelve *Cuscuta* species are marked in yellow ovals. Genome size and chromosome number of *Ipomoea* species are from previous studies (Hoshino *et al.*, 2016; Yang *et al.*, 2017). Ploidy level was determined based on a basic chromosome number (x) = 15 for monocentric *Cuscuta* and *Ipomoea* species, and x = 7 for holocentric *Cuscuta* species. Average chromosome size was calculated as the genome size divided by chromosome number and substantial differences in chromosome size are apparent in examples shown in the right-hand panel. Arrows mark primary constrictions. Chromosomes were counterstained with DAPI (blue). Scale bar represents 5 μm and is common to all chromosomes.

### Cuscuta karyotypes vary mainly due to polyploidization events and chromosome number reductions in holocentrics

In parallel to the genome size measurements, chromosome numbers and karyotypes were investigated using mitotic metaphase chromosome spreads prepared from shoot apical meristems (Fig. S1). Chromosome numbers (2n) varied from 14 to 60, and a majority of monocentric species had 30 or 32 chromosomes (Fig. 2). Additionally, three monocentric species in the subgenus *Grammica* (*C. gronovii*, *C. campestris*, and *C. pentagona*) had 2n = 60. The smallest chromosome numbers (2n = 14) were found in the two holocentric species *C. europaea* and *C. epithymum*, while *C. epilinum* had 2n = 42. Considering the chromosome numbers in monocentric *Cuscuta* species (Fig. 2) and in species belonging to related genera in Convolvulaceae, such as *Ipomoea* (Chen *et al.*, 2020), it is likely that the ancestral basic chromosome number (x) in the *Cuscuta* genus was 15. Consequently, we considered monocentric *Cuscuta* species either diploid (2n = 2x = 30 or 32) or tetraploid (2n = 4x = 60). In the three holocentric species, the basic chromosome number was probably reduced to seven; therefore, we identified *C. epithymum* and *C. europaea* as diploid (2n = 2x = 14) and *C. epilinum* as hexaploid (2n = 6x = 42). To test if the reduced number of chromosomes in this subgenus was due to chromosome fusions, we searched for possible relics of these events in the form of interstitial loci containing remnants of telomeric sequences. FISH using the telomeric probe (AAACCCT)6 revealed expected signals at chromosome termini in all tested species; however, interstitial loci were detected exclusively in the three species of the subgenus *Cuscuta* (Figs 3, S1). These results suggested that chromosome fusions indeed contributed to the reduction in chromosome counts in holocentric species.

**Figure 3.**
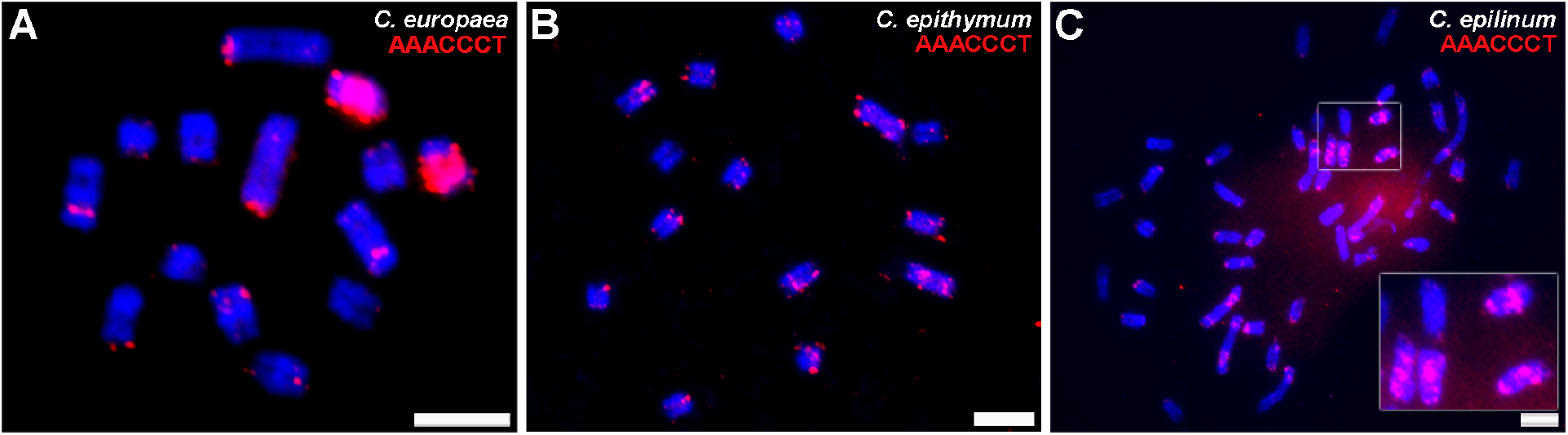
Detection of the telomeric sequence (AAACCCT)n in holocentric *Cuscuta* species. **(a)** *C. europaea,* **(b)** *C. epithymum,* and **(c)** *C. epilinum.* Note that some chromosomes in all three species display intercalary signals of the telomeric probe (red). These signals were in many cases considerably stronger compared with those at telomeres. Chromosomes were counterstained with DAPI (blue). All scale bars = 5 μm.

### Genome size variation is largely due to the amplification of repeats

Since polyploidization could explain only a small fraction of the genome size variation observed in *Cuscuta*, we investigated the contribution of the genome composition, especially the proportion of repetitive DNA. Low-pass whole genome sequencing was performed using the Illumina platform to generate 100 bp paired-end reads corresponding to 0.01–1.3 genome equivalents in all of the species. Highly and moderately repetitive sequences were then identified and characterized by graph-based clustering of sequence reads as implemented in RepeatExplorer (Novák *et al.*, 2013) and TAREAN (Novák *et al.*, 2017). All clusters representing at least 0.01% of the genome were manually checked, and their automated annotation was corrected if needed and finally used to characterize and quantify the most abundant repeats in each species. The fraction of low-copy repeats could not be evaluated by a clustering analysis owing to the extreme genome size variation among species. Therefore, we estimated the total proportion of repetitive (>20 copies/1C) and single/low-copy (≤20 copies/1C) sequences using all-to-all comparisons of forward Illumina reads representing 5% of the nuclear genome size. The global repeat composition of individual species is summarized in Fig. 4 and is listed in detail in Table S2. To evaluate the repeat content with respect to genome size differences between species, we expressed estimated genomic proportions of individual repeat types (Fig. 4a) as the total lengths per haploid genome (Fig. 4b, Table S2).

**Figure 4.**
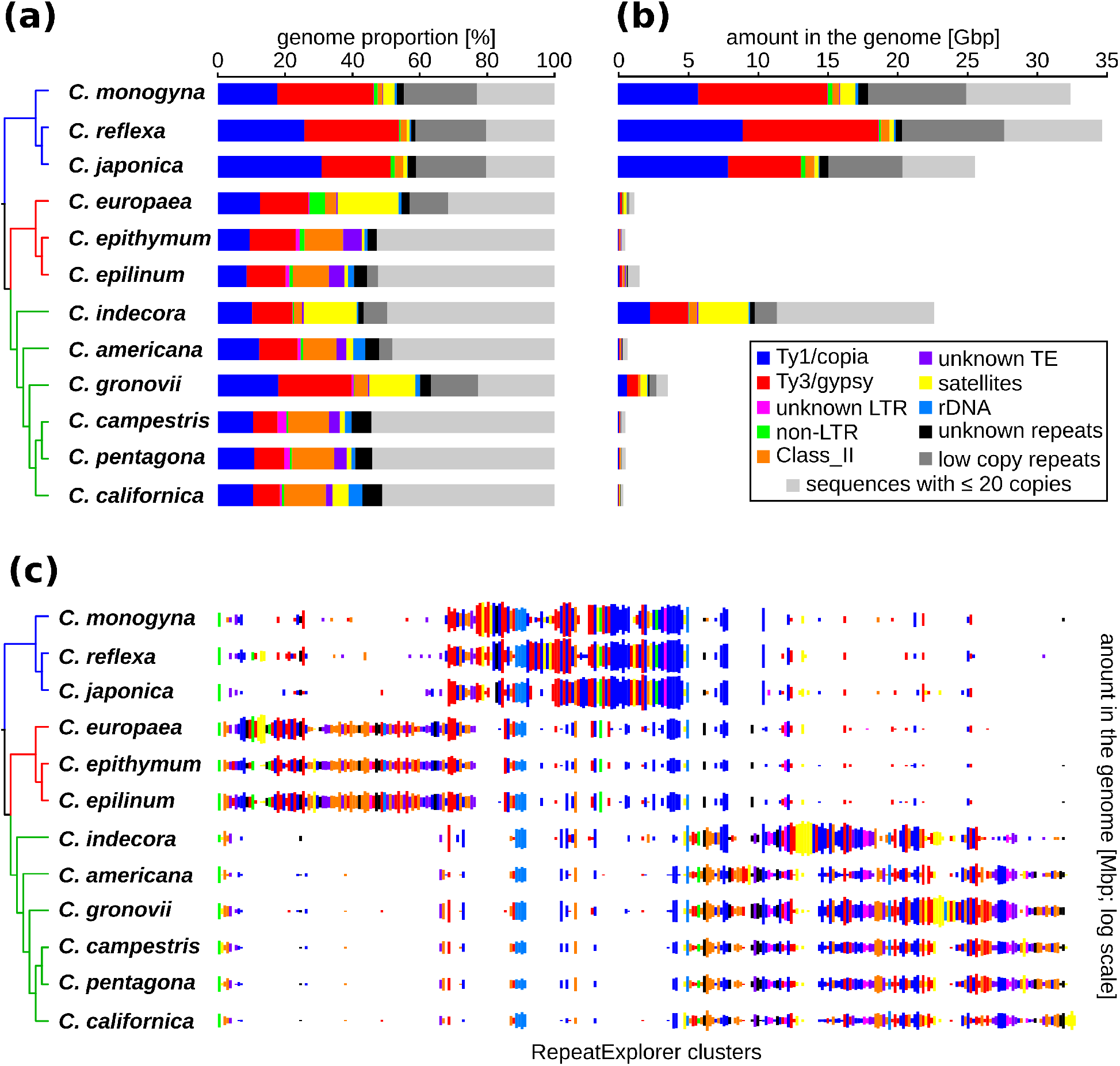
Composition of repetitive sequences in twelve *Cuscuta* species. Relative **(a)** and absolute **(b)** abundance of repetitive sequences in individual *Cuscuta* genomes as revealed using RepeatExplorer. **(c)** Comparison of repeat compositions among *Cuscuta* species inferred from a comparative RepeatExplorer analysis. The columns represent different RepeatExplorer clusters, (i.e., different repeats), and the size of the rectangles is proportional to the number of reads in a cluster for each species. Note that the composition of repetitive sequences is relatively similar in species of the same subgenus but differs considerably among the three subgenera.

LTR retrotransposons represented the major fraction of annotated repeats in all analyzed genomes. Although all plant LTR retrotransposon lineages (Neumann *et al.*, 2019) were detected, they differed in quantity, with SIREs being dominant among Ty1/copia and Tekay among Ty3/gypsy lineages (Table S2). Total genome proportions of LTR retrotransposon superfamilies ranged from 8.5% to 30.8% for Ty1/copia and 7.3% to 28.5% for Ty3/gypsy, and their ratios in individual genomes ranged from 1:0.67 in *C. japonica* to 1:1.61 in *C. monogyna.* Class II elements (DNA transposons) represented relatively large proportions of several small genomes (e.g., 12.6% in *C. pentagona)* and were relatively less abundant in large genomes. A substantial portion of some *Cuscuta* genomes was satellite DNA (satDNA), reaching 18.0% in *C. europaea.* The second largest proportion (15.9%) of satDNA was found in *C. indecora;* due to its genome size, this translated to the largest total amount of satDNA among investigated species, totaling 3.6 Gbp/1C (Fig. 4b, Table S2).

### Horizontally acquired retrotransposons occur in *Cuscuta* genomes but did not drive genome size expansion

To investigate repetitive DNA evolution within the genus in detail, we performed a comparative clustering analysis of repeats in different species. In particular, we combined the same numbers (430,615) of randomly sampled Illumina reads from each species in a single dataset for a comparative analysis using RepeatExplorer. Repeat clusters contained reads from orthologous repeat families in different species, leading to their identification and quantification (Fig. 4c). The repeat composition was similar among species belonging to the same subgenus, but differed considerably among the three subgenera. Additionally, the repeat profiles reflected patterns of phylogenetic divergence. For example, the profiles of *C. campestris* and *C. pentagona* were nearly identical, reflecting the close phylogenetic relationship between these two species. The profiles were also very similar between diploid *C. epithymum* and hexaploid *C. epilinum*, suggesting that the latter species originated from the polyploidization of *C. epithymum*-related ancestor(s) or *C. epithymum* itself.

An important observation was that genome size increases were not primarily driven by the amplification of LTR retrotransposons that were recently acquired by horizontal transfer. Such events would be revealed as species-specific clusters of abundant repeats that are amplified in one species but absent in closely-related species. However, no such cases were found among the top 306 clusters accounting for more than 0.01% of analyzed reads. Instead, species with large genomes, like *C. indecora* and *C. gronovii*, shared abundant repeat families with related taxa with small genomes, and genome expansion could be explained by the amplification of these shared repeats (Fig. 4c). Nevertheless, considering the parasitic strategy of *Cuscuta* species, we performed additional searches for cases of horizontal transfer (for details see Methods S2). First, we identified protein domain sequences that shared at least 80% similarity to non-Convolvulaceae sequences and exceeded the similarity to sequences from three Convolvulaceae species of the genus *Ipomoea* by at least 10% (Fig. S4a,b). Then, we compared full-length DNA sequences of the elements with sequences in GenBank. We found eight families of LTR retrotransposons that are not present in *Ipomoea* species but share very high similarity to elements from various non-Convolvulaceae species (Fig. S3c, Tables S7, S8). This indicates that these elements are either evolutionarily conserved and were selectively lost in some species or that they resulted from horizontal transfer. The occurrence of seven of these families in multiple *Cuscuta* species further suggests that horizontal transfer preceded the divergence of extant *Cuscuta* species. The proportion of these elements in *Cuscuta* genomes was relatively low, reaching a maximum of 0.32% in the case of the CRM family in *C. indecora.* Thus, their impact on *Cuscuta* genome size variation was very low.

### Holocentric species lack centromeric retrotransposons and vary in the proportions and chromosomal distribution of satDNA

Taking advantage of the phylogenetic position of holocentrics with respect to related monocentric species, we asked whether differences in repeat composition could be specifically linked to the transition to holocentricity. Disregarding the differences that could be simply attributed to variation in genome size, we found a qualitative pattern related to individual lineages of LTR retrotransposons. While all LTR retrotransposon lineages recognized in green plants (Neumann *et al.*, 2019) were detected in both subgenera of monocentrics, the holocentrics lacked CRM elements, a lineage of Ty3/gypsy chromoviruses known to preferentially insert into centromeres (Neumann *et al.*, 2011). To check whether CRM sequences could be missed in these species owing to their presence in the fraction of uncharacterized low-copy repeats, we searched contigs from the RepeatExplorer clusters representing these repeats using DANTE (Novák *et al.*, 2019). A single significant hit corresponding to the CRM protease domain was detected in a *C. epithymum* contig sequence estimated to occur in only two to three copies per holoploid genome. Such a low abundance or even the absence of CRMs in the other two holocentrics differed distinctly from the monocentric *Cuscuta* genomes, where these elements spanned at least 3–115 Mbp/1C (Table S2), which, considering the average element size of 6.8 kbp (Neumann *et al.*, 2019), corresponded to about 440–16,900 copies. The presence of the elements was also confirmed by FISH in the monocentric *C. gronovii*, where the CRM probe labeled centromeric regions on all chromosomes (Fig. 5a).

**Figure 5.**
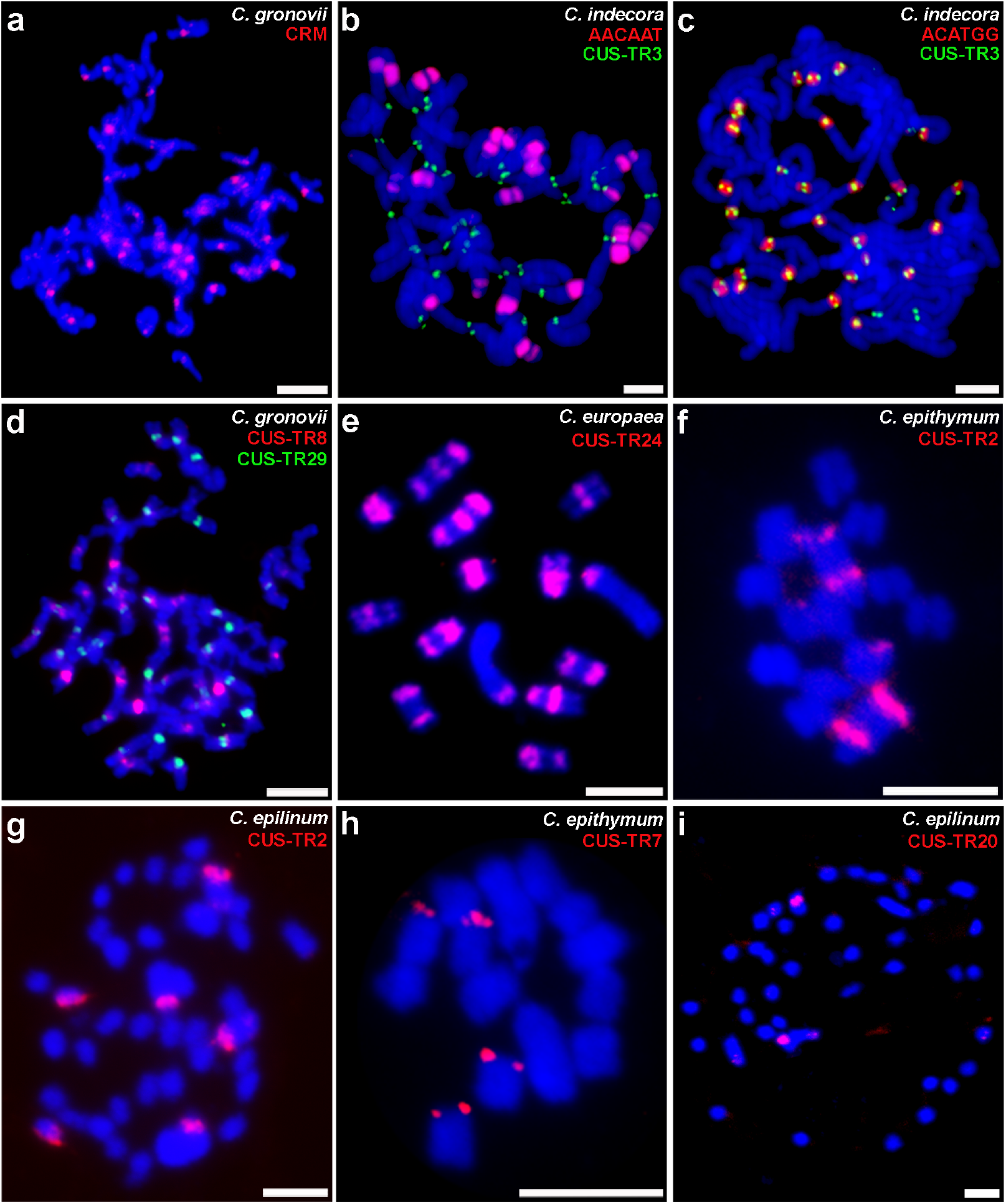
Detection of selected satDNA families and CRM retrotransposon on chromosomes of monocentric and holocentric *Cuscuta* species. **(a)** CRM retrotransposon (red) in *C. gronovii.* **(b)** AACAAT (red) and CUS-TR3 (green) in *C. indecora.* **(c)** ACATGG and CUS-TR3 (green) in *C. indecora.* **(d)** CUS-TR8 (red) and CUS-TR29 (green) in *C. gronovii.* **(e)** CUS-TR24 in *C. europaea.* **(f,g)** CUS-TR2 (red) in *C. epithymum* and *C. epilinum.* **(h)** CUS-TR7 (red) in *C. epithymum.* **(i)** CUS-TR20 (red) in *C. epilinum.* Chromosomes were counterstained with DAPI (blue). All scale bars = 5 μm.

SatDNA repeats are frequently associated with centromeres in plants, including holocentric species where they are located in longitudinal centromere grooves formed in each sister chromatid of metaphase chromosomes (Marques *et al.*, 2015). We identified 133 putative satDNA families that differed in abundance and distribution among *Cuscuta* species. Most (99) had monomers of tens to hundreds of nucleotides long, a typical size range for satDNA (Table S8); however, there were also 34 repeats with monomers shorter than 10 bp, corresponding to highly amplified, simple sequence repeat-like satellites (Table S9). They were particularly abundant in *C. indecora,* where FISH revealed that one of the most abundant SSR-like satellites, AACAAT, formed blocks of interstitial heterochromatin, while ACATGG was pericentromeric and flanked the putative centromeric satellite CUS-TR3 (Fig. 5b,c). The investigation of additional abundant satellites in the search for those with the centromeric localization revealed CUS-TR29 in *C. gronovii* (Fig. 5d). None of the satellites was shared by all species and only 12 were found in at least two species (Table S8), suggesting that satDNA exhibited fast turnover in *Cuscuta* species.

The three holocentric *Cuscuta* species differed considerably with respect to the satDNA content. *C. epithymum* and *epilinum* had low genomic proportions of satDNA (0.8% and 1.1%, respectively), whereas *C. europaea* had the highest genome proportion (18%) of satDNA among all analyzed species (Fig. 4, Table S2). Remarkably, the high proportion of satDNA in *C. europaea* was due to the massive amplification of a single satDNA family, CUS-TR24, which itself made up 15.5% of the genome (Table S8). FISH revealed that this repeat colocalized with a majority of the conspicuous heterochromatic bands (Fig. 5e and (Oliveira *et al.*, 2020)). We have recently shown that all other satellites in *C. europaea* occur at discrete loci (Oliveira *et al.*, 2020). In this study, we further revealed that CUS-TR2, the only satellite shared by all three holocentric species, also occurs at a few discrete sites in both *C. epithymum* and *C. epilinum* (Fig. 5f,g). Discrete localization was detected for CUS-TR7 and CUS-TR20 in *C. epithymum* and *C. epilinum,* respectively (Fig. 5h,i). The remaining two satellites in *C. epithymum* (CUS-TR22 and 23) were not detected. CUS-TR21 was detected at a single subtelomeric locus on one chromosome pair in *C. epilinum.* The chromosomal localization of other low-abundant satellites (≤0.4 Mbp; CUS-TR46, 47, and 82) in this species was not tested.

## Discussion

### Genome size variation among *Cuscuta* species is unusually high among plants

The genome size (1C) in *Cuscuta* species varies from 342 Mbp in *C. australis* to 34,734 Mbp in *C. reflexa* (Fig. 1, Table S6). This corresponds to a 102-fold difference, which is the largest variation documented for plant species in the same genus (Fig. S3a), similar in magnitude to that observed in entire plant families (Fig. S3c,d). Compared with the genus *Cuscuta*, which evolved 20.5–23 Ma (Fig. S2b), a number of plant families with substantial genome size variation among species emerged much earlier, including Melanthiaceae (65 Ma; (Vinnersten & Bremer, 2001), Santalaceae (>39±12 Ma, estimate for *Thesium;* (Moore *et al.*, 2010)), Liliaceae (85 Ma; (Kim & Kim, 2018)), Ranunculaceae (108–90 Ma; (Wang *et al.*, 2016)), Asparagaceae (66–48 Ma; (Raman *et al.*, 2019)).

On the other hand, the absolute size difference between the smallest and the largest *Cuscuta* genomes (i.e., 34,392 Mbp) is not as high as that in some other plant genera that include species with very large genomes, such as *Paris japonica* (Melanthiaceae; 1C = 149,185 Mbp). However, contrary to *C. australis* and *C. reflexa*, which are both diploids, most genera with greater variation in genome size either include polyploid species or species with unknown ploidy (Fig. S3b). The only known genus with greater absolute size difference among diploid species than that of *Cuscuta* is *Fritillaria,* in which the smallest genome of *F. davidii* (1C = 33,320 Mbp) and the largest diploid genome of *F. koidzumiana* (1C = 85,417 Mbp) differ by 52,097 Mbp.

### Impact of parasitism on genome size evolution

It has been proposed that parasitism leads to larger genome sizes because parasitic plants escape constraints imposed by root meristem growth rates (Gruner *et al.*, 2010) or simply due to the “genomic economy” in which resources are exploited from the host (Piednoël *et al.*, 2012). Indeed, in Orobanchaceae, obligate parasitic species have larger genomes than those of autotrophic and hemiparasitic species (Piednoël *et al.*, 2012). In addition, the hemiparasitic species *Viscum album* (Santalaceae) has one of the largest genomes recorded in plants (100,842 Mbp; (Zonneveld, 2010)). Other parasitic species in the family Santalaceae have much smaller genomes than those of *Viscum,* ranging from 255 to 6076 Mbp, and genome sizes of species from the parasitic family Loranthaceae are 2,842–17,150 Mbp (Pellicer & Leitch, 2020). These results suggest that the causality between parasitism and genome size increases may not be valid in general. Our data demonstrate that the parasitic lifestyle did not result in a genome size increase in *Cuscuta* species (Fig. 2). Relatively small genomes of other Convolvulaceae species included in the C-value database, including the genera *Convolvulus* (539–1686 Mbp/1C), *Ipomoea* (619–1,960 Mbp/1C), and *Calystegia* (696–784 Mbp/1C), suggest that the ancestral *Cuscuta* genome was also rather small and that genome expansion occurred later in the evolution of the genus in only a subset of species. Other species, such as *C. californica* and *C. australis*, likely exhibited genome shrinkage. The differences in genome size cannot be attributed to a different level of dependence on hosts because all *Cuscuta* species are obligate parasites and acquire all inorganic and almost all organic nutrients from their hosts. Most *Cuscuta* species retain photosynthetic ability, but it is greatly reduced compared with that in other plants and up to 4-fold differences in photosynthetic capacity do not correlate with phylogeny (Machado & Zetsche, 1990; Revill et al., 2005). Thus, it appears unlikely that the differences in the generally very low photosynthetic ability had a significant impact on genome size evolution in *Cuscuta*.

### Extent and causes of genome size variability differ among Cuscuta subgenera

Although polyploidy exists in *Cuscuta* species, all large genomes are diploid, indicating that the main driving force behind the genome size increase was the amplification of repetitive DNA (Fig. 4). The increase was particularly massive in the subgenus *Monogynella*, in which 1C genome sizes vary from 20,511 Mbp in *C. exaltata* (McNeal *et al.*, 2007a) to 34,734 Mbp in *C. reflexa*, and the three species included in this study contained 77–80% repetitive DNA (Figs 2, 4). Two other *Monogynella* species, *C. lupuliformis* and *C. monogyna*, also have large genomes (Pazy & Plitmann, 1995; McNeal *et al.*, 2007a), suggesting that all species in this subgenus evolved towards large genomes. By contrast, the subgenus *Cuscuta* lacks species with large genomes (Fig. 2 and (Pazy & Plitmann, 1995)), in agreement with the observation that holocentric species have small-to moderate-sized genomes (Bureš *et al.*, 2013). Although the genome size is only known for the three species included in this study, our data and the variation in chromosome number (Pazy & Plitmann, 1995; García, 2001; Guerra & García, 2004) suggest that genome size differences in this subgenus can be mostly attributed to polyploidy rather than to the amplification of repeats. The most highly variable subgenus in terms of both fold-change and absolute genome size difference is *Grammica*, in which 1C genome sizes of *C. australis* and *C. indecora* differ by 66-fold and 22,333 Mbp, although the two species diverged from a common ancestor only 10.6–8.4 Ma (Fig. S2b). The previously estimated genome size for *C. indecora* is even higher (32,115 Mbp/1C; (McNeal *et al.*, 2007a)); however, while <the *rbcL* sequence of *C. indecora* from the previous study (GenBank accession EU330274) was 99.8% to that in our study (Table S4), the ITS sequences (EU330311 and Table S3) shared only 89% similarity, suggesting that the two plants represented different species or subspecies. Unexpectedly, the proportion of repetitive DNA in *C. indecora* was only 50% (Fig. 4, Table S2), in sharp contrast to the large genomes of *Monogynella* species and to the *C. gronovii* genome (3,579 Mbp/1C) containing 77% repeats. Accounting for the large genome size and diploid number of chromosomes, we assume that a significant portion of single/low-copy sequences in the *C. indecora* genome were originally repetitive but diverged by mutation accumulation. In this scenario, which has been proposed for *Fritillaria* (Kelly *et al.*, 2015), the increase in genome size can be explained by a combined effect of repeat amplification and the infrequent removal of old copies. Although this is yet to be confirmed in *C. indecora*, our data indicate that the major drivers of genome size increase in this species differ from those in *C. gronovii* and *Monogynella* species.

### Horizontal transfer had a negligible impact on genome size evolution

Similar to many other plant species, a major part of repetitive DNA fraction in all *Cuscuta* species included in this study consisted of transposable elements, especially LTR retrotransposons (Fig. 4, Table S2; (Vitte & Panaud, 2005; Grover & Wendel, 2010; Macas *et al.*, 2015; Galindo-González *et al.*, 2017)). Although LTR retrotransposons are transmitted mainly vertically by the reproduction of host organisms, cases of horizontal transfer between species are relatively widespread in plants (Baidouri *et al.*, 2014). Since horizontal gene transfer is likely to be more frequent in parasitic plants and has already been proven in *C. reflexa* and *C. australis* (Zhang *et al.*, 2014; Yang *et al.*, 2016, 2019; Vogel *et al.*, 2018), it is conceivable that genome size differences among *Cuscuta* species could be due to the amplification of host-derived horizontally transferred elements. However, although we detected eight LTR retrotransposon families that may have been acquired via horizontal transfer, all but one were acquired before the divergence of the individual species in the genus and none of them achieved a high enough abundance to explain the dramatic differences in genome size. The strikingly lower similarity of many LTR retrotransposon RT domain sequences between *Cuscuta* and closely-related *Ipomoea* species than between *Cuscuta* and evolutionarily distant non-Convolvulaceae species (Fig. S4a) suggests that additional LTR retrotransposon families were acquired very early in the evolution of *Cuscuta*.

### Consequences of the transition to holocentricity in the subgenus *Cuscuta*

The transition to holocentricity in the subgenus *Cuscuta* was associated with substantial changes in the centromeric chromatin, repeat composition, and chromosome number. We have recently demonstrated in *C. europaea* that CENH3, a foundational kinetochore protein in most organisms regardless of centromere organization, either lost its function or acts in parallel to an additional CENH3-free mechanism for kinetochore positioning (Oliveira *et al.*, 2020). In this study, we found that H2AT120ph, thought to be a universal centromere marker in plants (Demidov *et al.*, 2014), is missing from the genomes of holocentric *Cuscuta* species.

These and perhaps additional yet unknown changes in centromeric chromatin may explain the absence of CRM retrotransposons in these species. CRM retrotransposons are preferentially localized in centromeres of a number of monocentric and some holocentric plant species (Neumann *et al.*, 2011; Marques *et al.*, 2015). Although the mechanism underlying their centromeric localization is not known, it may involve the specific recognition of centromeric chromatin during integration. Since CRM elements do occur in monocentric *Cuscuta* species (Table S2), we hypothesize that changes in centromeric chromatin in the subgenus *Cuscuta* prevented new insertions of CRM elements, and that old CRM insertions either gradually degenerated by accumulating mutations or were eliminated from the genomes via illegitimate recombination (Devos *et al.*, 2002). Contrary to most monocentric plant species and some holocentric species of the genus *Rhynchospora* (Plohl *et al.*, 2014; Marques *et al.*, 2015; Ribeiro *et al.*, 2017), in which centromeric chromatin is associated with satDNA, none of the satDNA families identified in holocentric *Cuscuta* species were distributed along entire chromosomes, as expected for centromere-specific sequences on holocentric chromosomes. No centromere-specific repeats have been found in holocentric *Luzula elegans*, which contains 20 satDNA families constituting 9.9% of the genome (Heckmann *et al.*, 2013). These satellites are enriched in centromere-free chromosome termini and at least some are involved in the end-to-end association of homologous chromosomes during meiosis (Heckmann *et al.*, 2014). Based on this observation, the presence of satDNA or heterochromatin in general may be required for proper behavior of holocentric chromosomes during meiosis. However, while the high amount and distribution of satDNA in *C. europaea* is in agreement with this notion (Oliveira *et al.*, 2020), the low genome abundance and rare occurrence of satDNA-containing loci on chromosomes in *C. epithymum* and *C. epilinum* (Fig. 5f-i, Table S8) suggest that satDNA is not necessary for the proper segregation of holocentric chromosomes.

The transition to holocentric chromosomes also led to a dramatic decrease in chromosome number in species in the subgenus *Cuscuta*. In the three species included in this study and previously studied species, the basic chromosome numbers are multiples of seven, including 2n = 2x = 14 (*C. europaea, C. epithymum* and *C. planiflora),* 2n = 4x = 28 (*C. approximata* and *C. palaestina)* and 2n = 6x = 42 (*C. epilinum* (Fig. S1b and (Pazy & Plitmann, 1991; García, 2001)). Chromosome numbers in other species of the subgenus are not multiples of seven, including 2n = 8, 10, 18, 20, 30, or 34 ((Pazy & Plitmann, 1995; García, 2001) and references therein). This variation indicates that multiple karyotype rearrangements followed the transition to holocentricity, consistent with the assumption that holocentric chromosomes have increased tolerance to chromosome fusions and fissions (Melters *et al.*, 2012).

## Acknowledgements

We thank Ms. Vlasta Tetourová and Ms. Jana Látalová for their excellent technical assistance. This research was financially supported by grants from the Czech Science Foundation (17-09750S) and the Czech Academy of Sciences (RVO:60077344). JČ and JD were supported by the ERDF project “Plants as a tool for sustainable global development” (No. CZ.02.1.01/0.0/0.0/16_019/0000827).

## Author contribution

PNe and JM conceived the study and designed the experiments. LO, T-SJ, KS, and SK performed the cytogenetics experiments. JČ and JD estimated nuclear genome sizes. AK performed DNA isolation and cloning experiments. PNe and PNo analyzed the sequence data. PNe and JM wrote the manuscript with input from LO, T-SJ, SK, KS, PNo, JČ, and JD. All authors read and approved the final manuscript.

## Methods S1: Estimation of genome size

Nuclear genome size was estimated as described previously (Doležel *et al.*, 2007). Briefly, 30 mg of fresh stem tissue from each *Cuscuta* accession and 10 mg of leaf tissue of a reference standard were chopped together in 500 μl of Otto I solution (Otto, 1990) using a sharp razor blade. The resulting homogenate was filtered through a 50 μm nylon mesh. Nuclei were then pelleted (300 × *g,* 3 min) and resuspended in 300 μl of Otto I solution. After 30 min of incubation on ice, 600 μl of Otto II solution supplemented with 50 μg/ml RNase and 50 μg/ml propidium iodide was added. Samples were analyzed using a CyFlow Space flow cytometer (Sysmex Partec GmbH, Görlitz, Germany) equipped with a 532 nm green laser. The gain of the instrument was adjusted so that the peak representing G1 nuclei of the standard was positioned approximately on channel 100 on a histogram of relative fluorescence intensity when using a 512-channel scale. Owing to the huge range of genome sizes, several internal reference standards were used for genome size estimation. *Raphanus sativus* cv. Saxa (1.11 pg/2C; (Doležel *et al.*, 1998)) was used as the internal reference standard for *C. californica* and *C. australis; Lycopersicon esculentum* cv. Stupické polní tyčkové rané (1.96 pg/2C; (Doležel *et al.*, 1992)) was used as the internal reference standard for *C. pentagona, C. campestris, C. americana, C. europaea* and *C. epithymum; Zea mays* cv. CE-777 (5.43 pg/2C; (Lysák & Doležel, 1998)) was used as the internal reference standard for *C. epilinum; Pisum sativum* cv. Ctirad (9.09 pg/2C; (Doležel *et al.*, 1998)) was used as the internal reference standard for *C. gronovii* and *Vicia faba* ssp. *faba* var. *equina* cv. Inovec (26.9 pg/2C; (Doležel *et al.*, 1992)) served as the internal reference standard for *C. indecora* and *C. japonica. Cuscuta japonica* was used as the internal reference standard for *C. monogyna* and *C. reflexa.* Three individual plants per each *Cuscuta* species were sampled, and each sample was analyzed three times, each time on a different day. A minimum of 5000 nuclei per sample were analyzed and 2C DNA contents (in pg) were calculated from the means of the G1 peak positions by applying the formula: 2C nuclear DNA content = (sample G1 peak mean) × (standard 2C DNA content) / (standard G1 peak mean). The mean nuclear DNA content (2C) was then calculated for each species. DNA contents in pg were converted to genome size in bp using the conversion factor 1 pg DNA = 0.978 Gbp (Doležel *et al*., 2003).

## Methods S2: *Identification of horizontally transferred LTR retrotransposons*

Possible cases of horizontally transferred LTR retrotransposons were identified by an analysis of sequences encoding conserved protein domains of LTR retrotransposons extracted from contigs assembled from the repeat clusters identified in individual species. These protein sequences were used for similarity searches against sequences from three *Ipomoea* species which belong to Convolvulaceae and are the most closely-related taxa to *Cuscuta* with available genome sequence data. The sequence data for *Ipomoea* species included genome assemblies of *I. batatas, I. nil,* and *I. trifida* (GenBank assembly accessions GCA_002525835.2, GCF_001879475.1, and GCA_000978395.1; (Hirakawa *et al.*, 2015; Hoshino *et al.*, 2016; Yang *et al.*, 2017) and RepeatExplorer contigs generated for the two former species using Illumina sequence data downloaded from SRA (run accessions ERR1512971 and DRR013919). The same *Cuscuta* protein sequences were compared with REXdb, a database of retrotransposon protein domains retrieved from a wide range of green plant taxa but lacking sequence entries from Convolvulaceae (Neumann *et al.*, 2019). In theory, vertically inherited retrotransposon sequences in *Cuscuta* should generate the best similarity hits to corresponding elements from *Ipomoea* genomes due to the close phylogenetic relationship of the two genera. Detecting higher similarity to sequences from phylogenetically distant taxa would imply that horizontal transfer occurred. Thus, sequences that displayed at least 80% similarity to non-Convolvulaceae sequences from REXdb and exceeded the similarity to *Ipomoea* sequences by at least 10% were selected for further analysis. Representative full-length element DNA sequences of the selected elements were either extracted from the genome assembly of *C. campestris* (Vogel *et al.*, 2018) or were manually reconstructed using RepeatExplorer. The absence of related sequences in the three *Ipomoea* species and their presence in non-Convolvulaceae species was confirmed by BLASTN searches.

## Notes S1: Origin of *Cuscuta* species

Seeds of *C. americana* L. (serial number: 0513111), *C. californica* Choisy (serial number: 0496988), *C. europaea* L. (serial number: 0101147), *C. indecora* Choisy (serial number: 0532275) and *C. pentagona* (serial number: 0484019) were obtained from the Royal Botanic Garden (Ardingly, UK). Seeds of *C. campestris* Yunck., *C. gronovii* Willd. and *C. japonica* Choisy were provided by Dr. Chnar Fathoulla (University of Salahaddin, Kurdistan Region, Iraq), Dr. Mihai Costea (Wilfrid Laurier University, Waterloo, Ontario, Canada) and Dr. Takeshi Furuhashi (RIKEN Center for Sustainable Resource Science, Yokohama, Japan), respectively. Seeds of *C. epilinum* Weihe and *C. monogyna* Vahl. were obtained from the Leibniz Institute of Plant Genetics and Crop Plant Research (Gatersleben, Germany) and Lebanese Agricultural Research Institute (Rayak, Lebanon), respectively. *C. reflexa* Roxb. plant was obtained from the Botanic Gardens of the Rhenish Friedrich-Wilhelm University (Bonn, Germany). *C. epithymum* L. plants were collected from a natural population at “U Cáby” (Kroclov, Czech Republic). Seeds of *C. australis* Hook.f. were provided by Prof. Jianqiang Wu (Kunming Institute of Botany, Chinese Academy of Sciences, Kunming, China).

**Figure S1.**
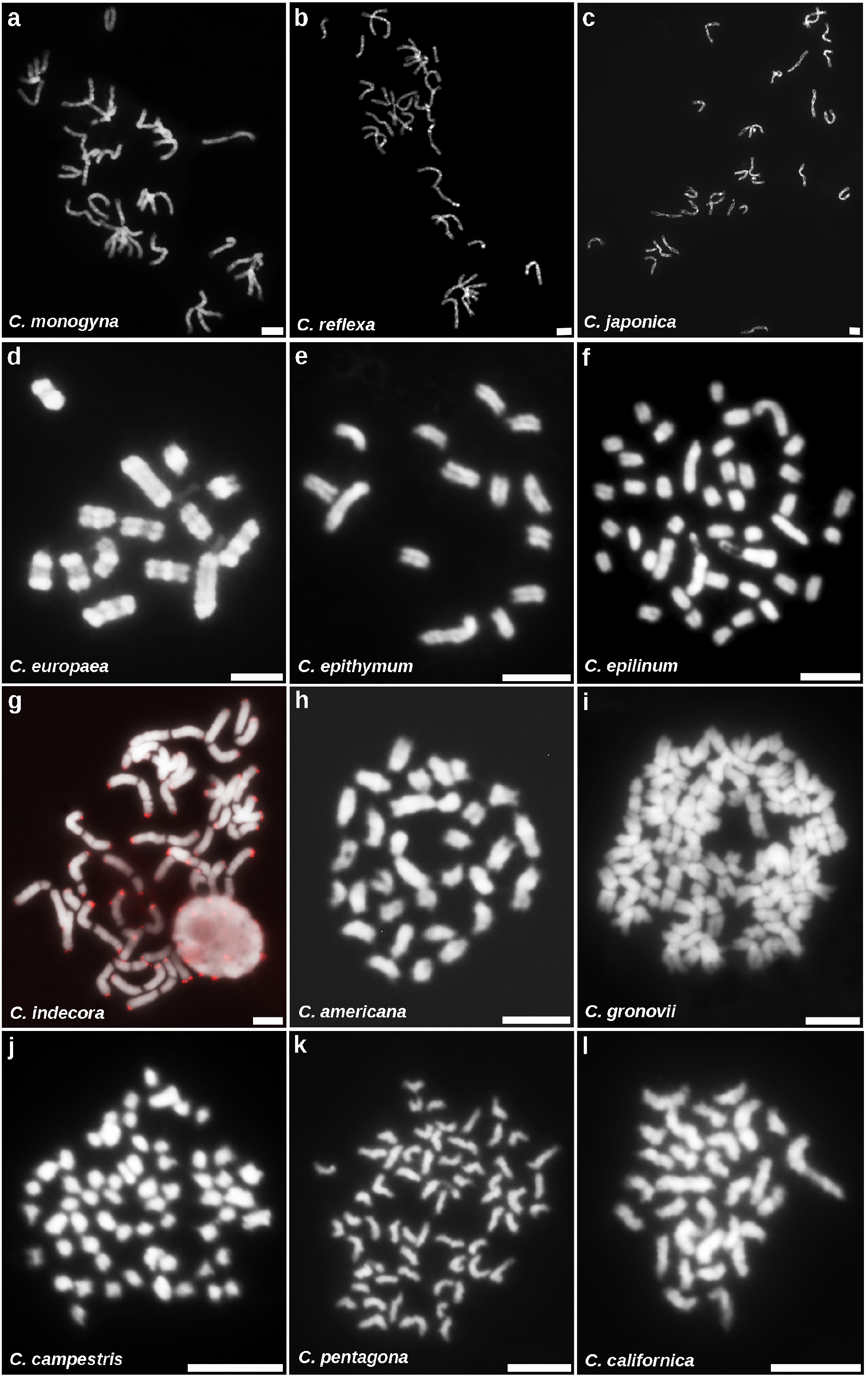
Mitotic metaphase chromosomes of all twelve *Cuscuta* species included in the study. The red signals on chromosomes of *C. indecora* show telomeric loci detected by FISH and helped to distinguish chromosome termini from primary and secondary constrictions that often resembled gaps. Chromosomes were counterstained with DAPI (blue). All bars = 5 μm.

**Figure S2.**
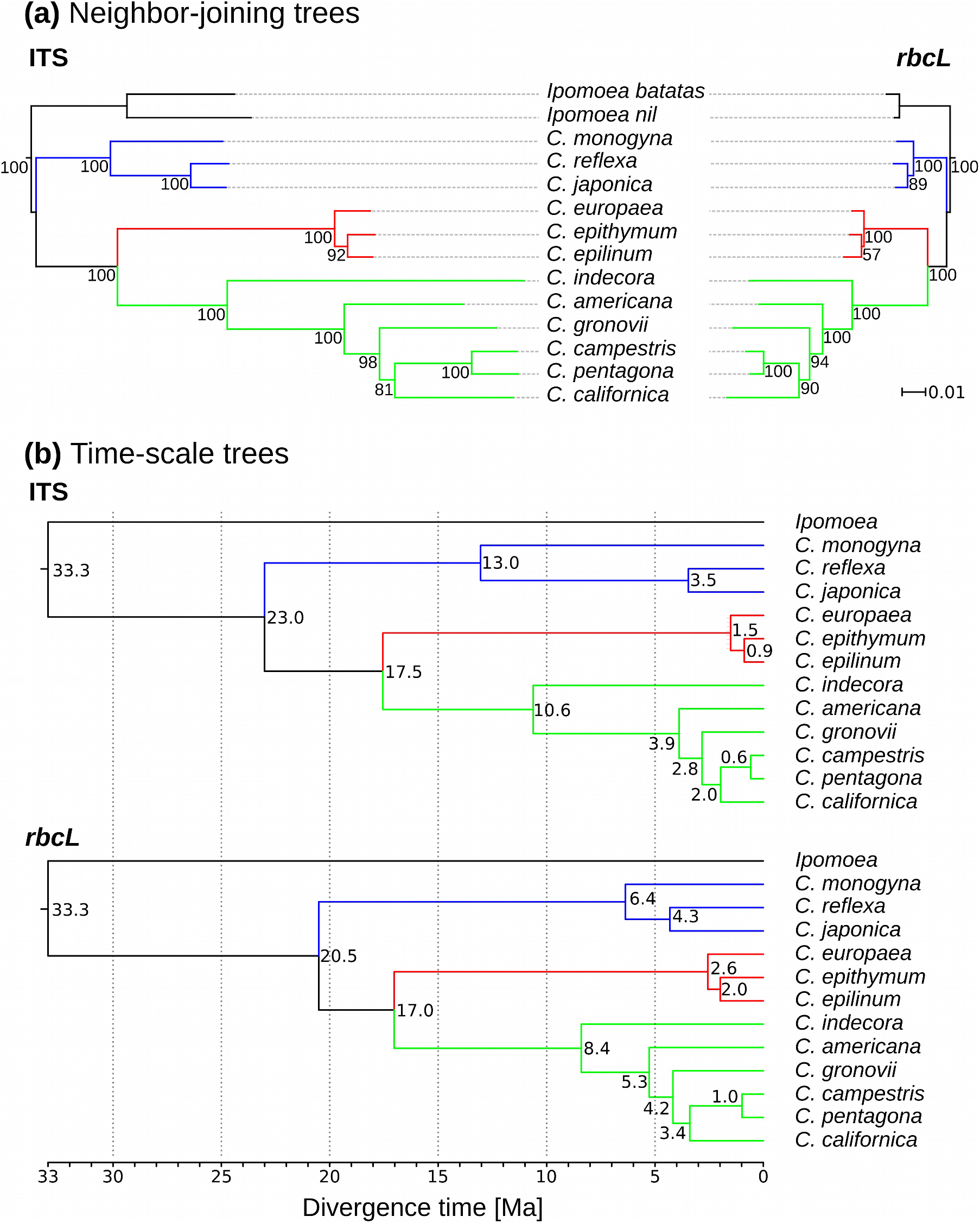
Phylogenetic trees inferred from ITS and *rbcL* sequence alignments. **(a)** Comparison of neighbor-joining trees inferred from alignments of ITS and *rbcL* sequences. The numbers at nodes are bootstrap values calculated from 5000 replicates. **(b)** Time-scale trees inferred using the maximum-likelihood method with smart model selection (Guindon *et al.*, 2010; Lefort *et al.*, 2017) and then dated using the RelTime method implemented in MEGA X (Mello, 2018) assuming that the most recent common ancestor of *Cuscuta* and *Ipomoea* existed 33.3 million years ago (Sun *et al.*, 2018). Branches corresponding to subgenera *Monogynella, Cuscuta* and *Grammica* are highlighted in blue, red and green, respectively.

**Figure S3.**
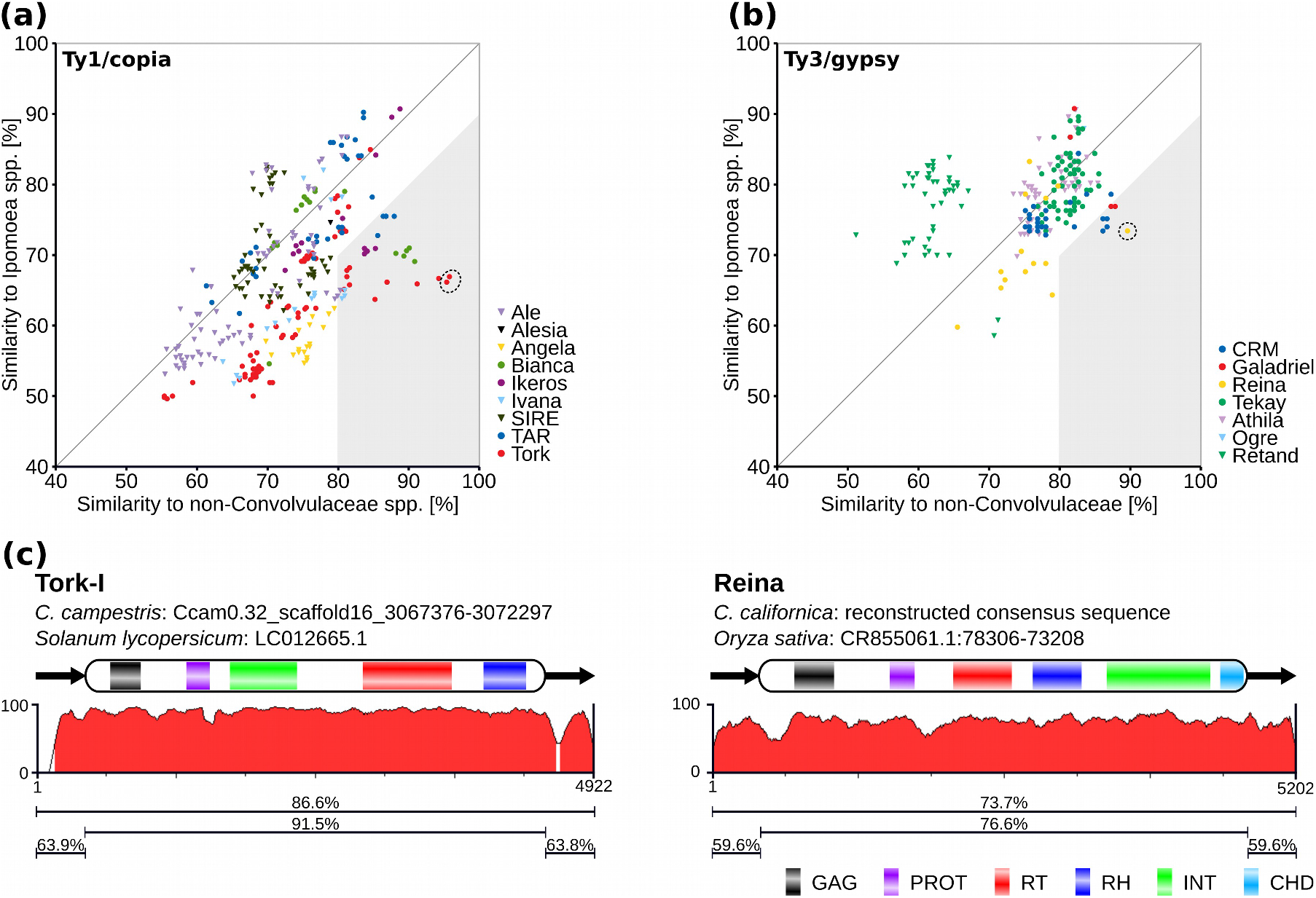
Detection of putative horizontally transferred LTR retrotransposons. **(a,b)** Plots of highest similarities between *Cuscuta* Ty1/copia **(a)** and Ty3/gypsy **(b)** RT domain protein sequences and *Ipomoea* and non-Convolvulaceae sequences from REXdb (Neumann *et al.*, 2019). Despite the expectation that *Cuscuta* sequences would be more similar to sequences from closely-related *Ipomoea* species, many sequences, particularly Ty1/copia elements, shared higher similarity with sequences from various non-Convolvulaceae species. Dots in the shaded area represent sequences that displayed at least 80% similarity to non-Convolvulaceae sequences and exceeded the similarity to *Ipomoea* sequences by at least 10%. These sequences were selected for further analysis as candidate horizontally transferred elements. **(c)** Examples of two retrotransposon families predicted to have been horizontally transferred and similarity profiles produced using zPicture (Ovcharenko *et al.*, 2004). RT domains of these elements are marked in circles in the similarity plots.

**Figure S4.**
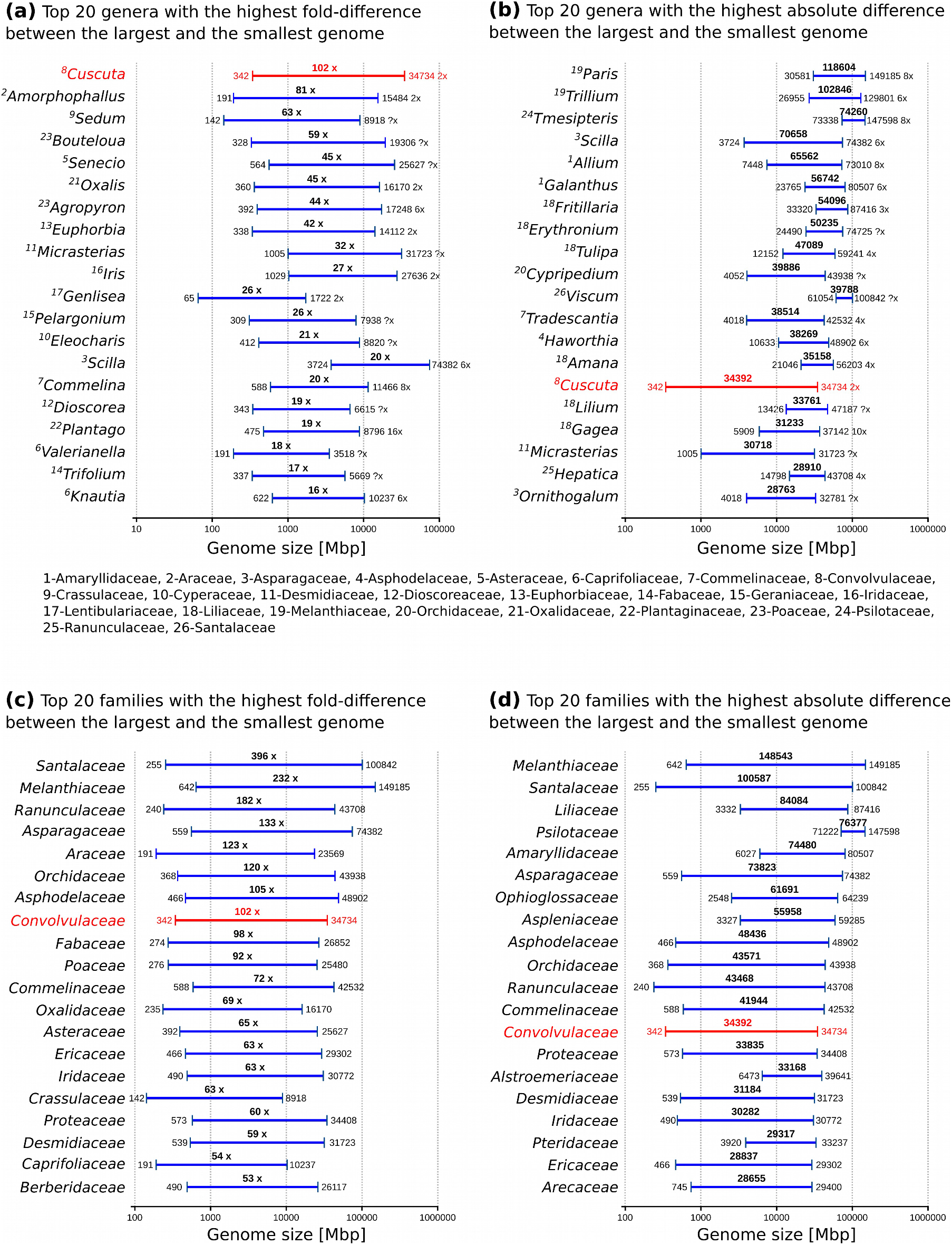
Comparison of genome size variation in *Cuscuta* and other groups of species with high genome size variation. Comparisons were performed at the genus **(a,b)** and family **(c,d)** levels. Data were acquired from the plant DNA C-values database (release 7.1; (Pellicer & Leitch, 2020)). Only prime genome size estimates were included in the comparison. Numbers above the lines correspond to fold (a,c) or absolute (b,d) differences between the smallest and largest genomes. The genus *Cuscuta* and family Convolvulaceae are highlighted in red. Numbers following the maximum genome size in each genus specify the ploidy level (x) of the respective species (question mark indicates unknown ploidy).

